# Fluctuation of lysosomal protein degradation in neural stem cells of postnatal mouse brain

**DOI:** 10.1101/2023.05.12.540513

**Authors:** He Zhang, Karan Ishii, Tatsuya Shibata, Shunsuke Ishii, Marika Hirao, Zhou Lu, Risa Takamura, Satsuki Kitano, Hitoshi Miyachi, Ryoichiro Kageyama, Eisuke Itakura, Taeko Kobayashi

## Abstract

Lysosomes are intracellular organelles responsible for degrading diverse macromolecules delivered from several pathways, such as the endo-lysosomal and autophagic pathways. Recent reports have suggested that lysosomes are essential in regulating neural stem cells in developing, adult, and aged brains. However, the activity of these lysosomes has not yet been monitored in these brain tissues. Here, we report a new probe to measure lysosomal protein degradation in brain tissue by immunostaining. Our results demonstrate the fluctuation of lysosomal protein degradation in neural stem cells depending on age and brain disorder. Neural stem cells increase lysosomal activity during hippocampal development in the dentate gyrus, but aging and aging-related disease reduces their activity. In addition, physical exercise increases lysosomal activity in neural stem cells and astrocytes. We hypothesize three different stages of lysosomal activity: the increase in development, the stable state for the adult stage, and the reduction by damages with age or disease.

---

Lysosomes are membrane-bound organelles for the degradation of biological macromolecules and for a hub of signal transduction after sensing intracellular amino acid levels (Ballabio & Bonifacino, 2020). They also function for lipid metabolism and calcium storage. Recent reports suggested that lysosomal regulations play essential roles in neural stem cells (NSCs) in developing, adult, and aged brains (Kobayashi *et al*, 2019; Leeman *et al*, 2018; Yuizumi *et al*, 2021). In the developmental stage, neural stem/progenitor cells (NSPCs) contain much more lysosomes than neurons of the developing telencephalon (Yuizumi *et al*., 2021). The deficiency leads to premature differentiation of NSPCs via lowered expression of the lysosomal transporter for histidine and peptides (Yuizumi *et al*., 2021). Lysosomes are more enriched in slowly dividing NSPCs, the likely source of adult NSCs. Enhanced lysosomal biogenesis by forced expression of constitutively active mutants of TFEB, a master transcription factor for lysosomal biogenesis, suppresses the cell cycle of NSPC in the embryonic brain (Yuizumi *et al*., 2021). NSPCs enter the quiescent state to maintain NSCs for a long time in the adult brain. Quiescent NSCs contain enriched lysosomes, and their deficiency reactivates quiescent NSCs after the accumulation of activated EGF and Notch receptors (Kobayashi *et al*., 2019). Lysosomes in quiescent NSCs store protein aggregates (Leeman *et al*., 2018), and quiescence exit clears these aggregates to recover their proteostasis for proliferation (Morrow *et al*, 2020). Especially in the aged brain, quiescent NSCs accumulated more protein aggregates in their lysosomes, reducing reactivation ability. The enhancement of lysosomal biogenesis reduces their aggregates and recovers the reactivation from quiescent NSCs (Leeman *et al*., 2018). All of these reports demonstrate that lysosomes are involved in NSC maintenance in the brain; however, how the protein degradation by lysosomes alters at different stages and conditions in these NSCs *in vivo* has not yet been analyzed in detail. Recently, fluorescent probes have become available to monitor biological activities, including signal transduction and protein degradation (Mizushima & Murphy, 2020; Neefjes & Dantuma, 2004). In the adult brain, autophagy flux probes revealed that autophagy-lysosome function is impaired in neurons before pathogenesis with lowered v-ATPase activity (Lee *et al*, 2022). In this report, we show a novel lysosomal probe containing two fluorescent proteins of different stabilities to monitor lysosomal protein degradation activity in NSCs of the mouse brain from juvenile to old ages (Ishii *et al*, 2019; Yanai & Endo, 2021). Quantification by the lysosomal probe revealed that lysosomal activity in hippocampal NSCs 1) increased from juvenile to adolescence while 2) it was maintained in adults but was 3) finally damaged with age and pathology of Alzheimer’s disease (Oakley *et al*, 2006). These three different states of lysosomal degradation activity suggest the different roles of lysosomes in NSC maintenance in the brain.

## Lysosome probe to monitor protein degradation in lysosomes

To monitor lysosomal activity at single lysosome resolution *in vivo*, we generated a new variant of the Lysosomal-METRIQ (Measurement of protein Transporting integrity by RatIo Quantification) probe (Ishii *et al*., 2019), consisting lysosomal deoxyribonuclease (DNase) II alpha, and tandem fusion of two fluorescent proteins, mCherry and super-folder green fluorescent protein (sfGFP) (Khmelinskii *et al*, 2012). DNase II alpha is a lysosomal enzyme which is synthesized in the endoplasmic reticulum (ER) and transported to lysosomes through the Golgi apparatus (Ohkouchi *et al*, 2013). This lysosomal probe, named LysoMonitor (LyMo), is the tandem-fusion protein with short linker peptides between DNase II alpha and two fluorescent proteins (Fig. 1A), which have different stabilities against low pH and lysosomal proteases: mCherry is stable (half-life > 1 day), while sfGFP is unstable in lysosomes (Katayama *et al*, 2008; Pedelacq *et al*, 2006). LyMo signals in NSCs were compared by immunostaining with antibodies. Lysosomal marker Lamp1 completely colocalized with mCherry signals (Fig. 1B), and GFP signals were diffusively distributed in cells and colocalized with weaker mCherry signals than those in lysosomes because of including transporting LyMo through the ER and Golgi into lysosomes. This suggests that, compared to mCherry, the GFP region of LyMo is unstable and degraded rapidly after transport into lysosomes, regions with strong mCherry signals. The intensities of the GFP and mCherry signals were measured in thresholded mCherry “dots” of lysosomes. Then the lysosomal activity was calculated by dividing the GFP intensity by the mCherry intensity in each dot (Fig. 1C). The lysosomal inhibitor bafilomycin A1 (BafA) increased the GFP to mCherry signal ratio, while the mTORC1 kinase inhibitor Torin1 (Settembre *et al*, 2012) activated lysosomes through TFEB activation and decreased the GFP to mCherry signal ratio (Fig. 1B, C). These results confirmed that the ratio of GFP to mCherry signal is inversely correlated with lysosomal activity (Fig. 1C). To confirm this probe monitoring protein degradation in lysosomes, the stabilities of GFP and mCherry proteins were examined by cycloheximide-chase experiments and Western blotting. It revealed that 1) the GFP and mCherry fusion components were excised from the full length of the LyMo (upper bands around 50 kDa (mCherry-GFP fusion) and around 100 kDa (full length) in Fig. 1D), 2) the excised mCherry was stable, and 3) the excised GFP was rapidly degraded (mCherry and GFP bands at ∼25 kDa in Fig. 1D). GFP degradation depended on lysosomal activity and was inhibited by BafA (Fig. 1D, E). These results indicate that the calculated ratio of LyMo signals detected by immunostaining of GFP and mCherry reflects proteolytic activity in the lysosome.

**Figure 1.**
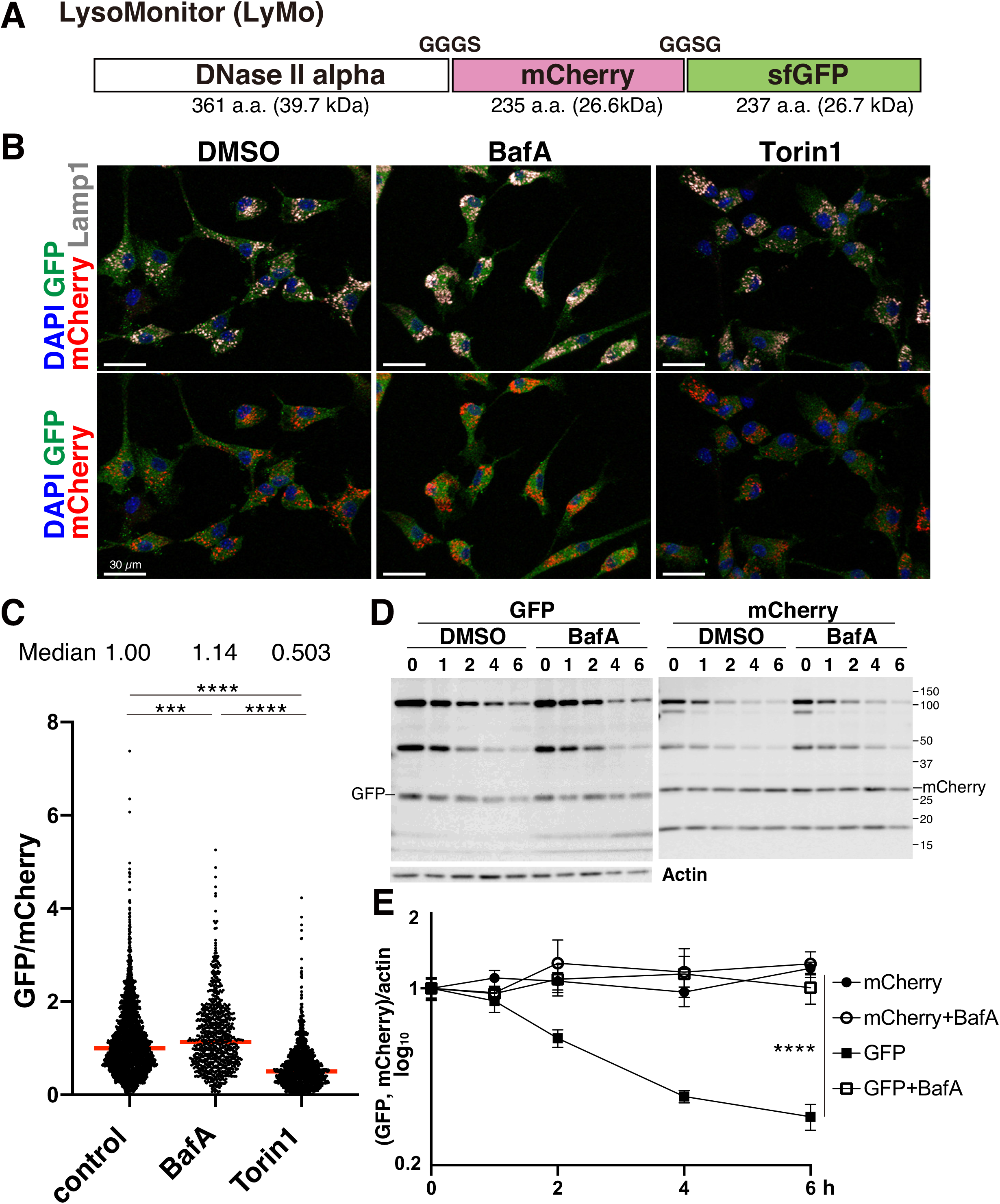
Novel lysosomal probe to monitor protein degradation in lysosomes. **A. Lysosome monitoring probe (LysoMonitor, LyMo)**. A fusion protein of lysosomal enzyme DNase II alpha, mCherry, and sfGFP with linker amino acids. **B. LyMo immunofluorescence imaging**. NSCs expressing the lysosome probe were fixed and immunostained for GFP (green), mCherry (red), and Lamp1 (grey) with DAPI (blue). NSCs were cultured with doxycycline for one day to induce LyMo expression by a TET-on system and incubated with DMSO, 20 nM bafilomycin A1(BafA), or 100 nM Torin1 for 4 h before fixation. Scale bars, 30 μm. **C. Quantification of LyMo**. GFP intensity divided by mCherry intensity was measured in mCherry-positive individual lysosomes and normalized with the control sample median. Red bars represent medians. (\*\*\**P* < 0.001, \*\*\*\**P* < 0.0001; one-way ANOVA with Tukey’s multiple comparisons tests). Median values are above the chart. **D, E. Protein stability of LyMo**. NSCs were cultured with doxycycline for one day and then incubated with 10 μg/ml cycloheximide with or without 20 nM bafilomycin A1 (BafA). Full-length LyMo and mCherry-sfGFP fusion proteins were commonly detected by GFP and mCherry antibodies around 100 and 50 kDa, respectively (D). GFP and mCherry bands around 25 kDa (noted as GFP and mCherry in Panel (D)) were measured and plotted in the chart after normalization with actin (E). Truncated mCherry (noted as mCherry* around 20 kDa) was not used for measurement. Data represent means ± s.e.m. (\*\*\*\**P* < 0.0001; two-way ANOVA with Tukey’s multiple comparison tests, n=4).

### LyMo transgenic mice to monitor lysosomal activity *in vivo*

LyMo was then applied to monitor lysosomal activity by immunostaining *in vivo* using transgenic (Tg) mice. To express LyMo in NSCs of the mouse brain, we generated Tg mice encoding LyMo under mouse glial fibrillary acidic protein (GFAP) promoter (Fig. 2A), which is an active promoter in adult NSCs and astrocytes (Seki *et al*, 2014). In the hippocampal dentate gyrus (DG), mCherry signals of LyMo were present in the granular cell layer and colocalized with Lamp1 (Fig. 2B, right panels). These mCherry dots were present in cells that were positive for NSC markers: Sox2, GFAP, and Nestin (Fig. 2B, left and center panels). To confirm these LyMo signals are available to quantify lysosomal activity in brain tissue, brain slices of the DG were immunostained with GFP and mCherry antibodies with DAPI after slice culture with lysosomal inhibitor, BafA (Fig. 2C). Measurement of GFP and mCherry intensities in the DG confirmed an increased ratio of GFP to mCherry in the presence of lysosomal inhibitor BafA (Fig. 2D). From this, we deduced that the LyMo quantification method by immunohistochemistry can accurately monitor relative lysosomal activity in tissue. Using this LyMo mouse, we investigated lysosomal activity in NSCs of the adult mouse brain after their activation by voluntary running since physical exercise has been shown to activate neurogenesis in the hippocampus (Garrett *et al*, 2012; van Praag *et al*, 1999). Transgenic young adult mice expressing LyMo were divided into individual cages either with or without free access to a running wheel and fixed after six weeks. Immunohistochemistry detected both an increase of approximately 2% of Ki-67-positive cells in Sox2-positive cells (from 13.8% to 15.9%) and an increase of newly-born neurons that were double positive for Ki-67 and DCX (from 16.4% to 22.7%), an immature neuron marker, in the DG of Tg mice in the wheel cages (Fig. 3A, B). These results are consistent with previous reports suggesting that physical exercise increased the efficiency of the production of newly-born neurons in the adult hippocampus. Next, we analyzed lysosomal activity in NSCs. Since the GFAP promoter is active in both NSCs and astrocytes, we immunostained with an S100β, astrocyte marker antibody to exclude the LyMo signals from astrocytes for measurement. S100β-positive cells were present in the DG even if LyMo-positive cells have a radial glia-like morphology, a typical morphology of NSCs in the DG (Fig. 3C, lower panels) (Karpf *et al*, 2022; Kriegstein & Alvarez-Buylla, 2009). LyMo signals in S100β-negative cells were quantified to calculate lysosomal activity in NSCs in the DG. Quantification revealed increased lysosomal activity in the hippocampal DG of mice in wheel cages for both NSCs and astrocytes because GFP/mCherry values in running mice were lower than those in controls (Fig. 3D). These results indicate that exercise enhances neurogenesis in the hippocampus, which is accompanied by an increase in lysosomal activity in both NSCs and astrocytes. Lysosomal activation in astrocytes and NSCs implies that the global changes occurred in the brain, such as previously reported beneficial effects on vascular systems, which are severely damaged with the aging process in the brain (Cole *et al*, 2022) or global improvements of metabolic systems. The results are consistent with the findings wherein exercise increases the expression of lysosomal-related proteins in the brain after running through TFEB activation *in vivo* (Huang *et al*, 2019). Increased lysosomal activity in both NSCs and astrocytes implies that an enhanced vascular function by physical exercise alters the metabolic state and increases overall lysosomal activity (Leiter *et al*, 2019).

**Figure 2.**
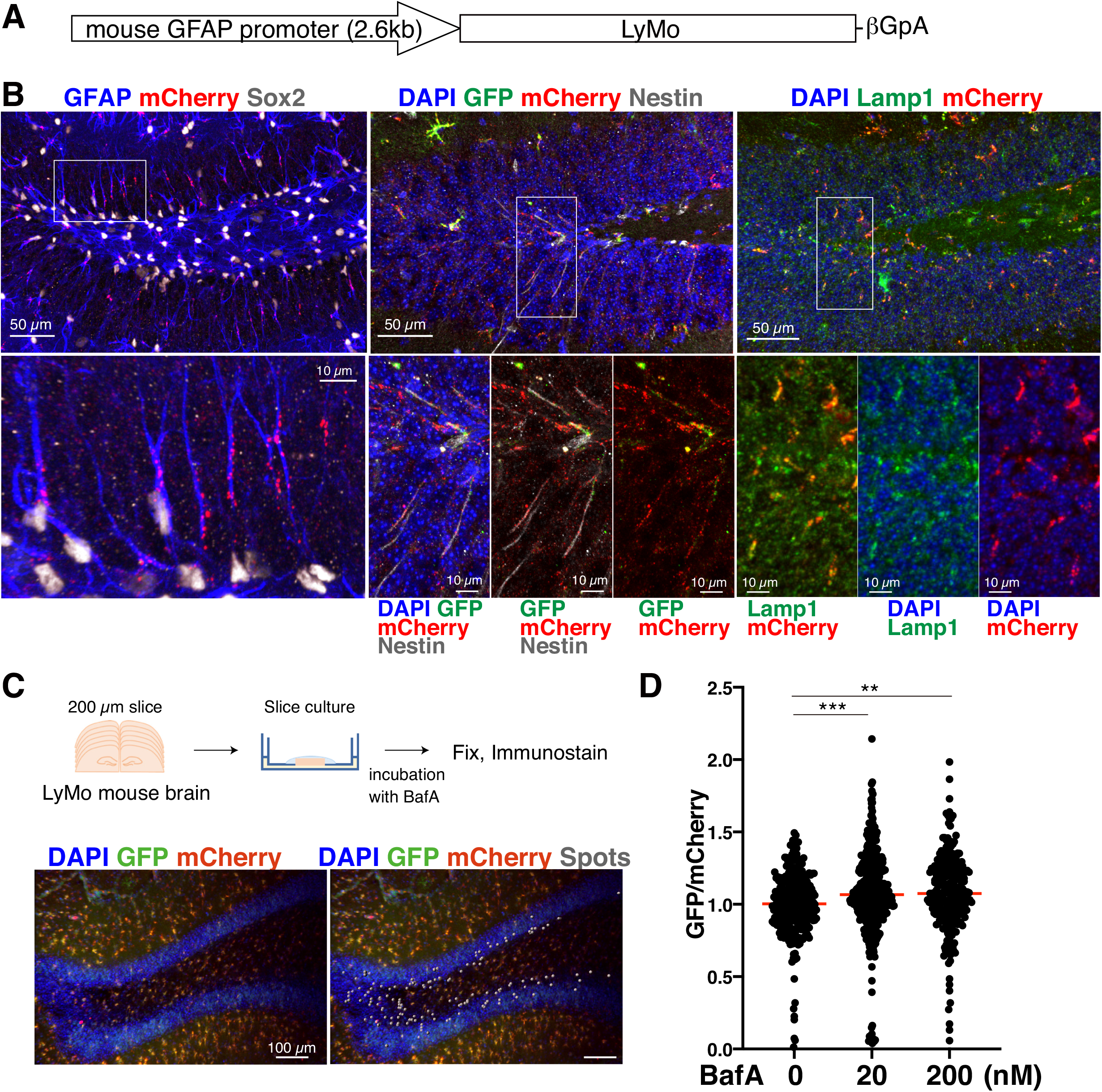
Transgenic mice to monitor lysosomal degradation activity in NSCs of brain tissue. **A. Schematic representation of LyMo construct for transgenic mice**. Mouse GFAP promoter drives LyMo protein expression. **B. Expression of LyMo in the hippocampal dentate gyrus**. Left: GFAP (blue) and Sox2 (grey)-positive NSCs with radial fiber express LyMo (mCherry, red). Middle: Nestin-positive NSCs express LyMo (GFP and mCherry, green and red). Right: LyMo expression (mCherry) colocalizes with Lamp1 (green). The area enclosed by the white squares in the upper panels was enlarged in the lower panels. No fluorescent signal was detected without IHC (not shown). **C. Slice culture of the LyMo mouse brain**. Upper: Schematic representation of the experimental flow of slice culture. 200 μm brain slices from LyMo mouse at 14 months old were cultured with or without bafilomycin A1(BafA) for one day and fixed for IHC. Lower: representative image of IHC (left) and measurement (right). Spots (grey) were put on mCherry dots in the granule cell layer (DAPI) to measure mCherry and GFP intensities in the dentate gyrus NSCs. **D. Quantification of LyMo in tissue slice**. GFP and mCherry ratio was measured in mCherry dots and normalized with the median of the control sample. Red bars represent medians. The ratio of GFP to mCherry was inversely correlated with lysosomal activity in tissue NSCs. (\*\**P* < 0.01, \*\*\**P* < 0.001; one-way ANOVA with Tukey’s multiple comparisons tests, n=3).

**Figure 3.**
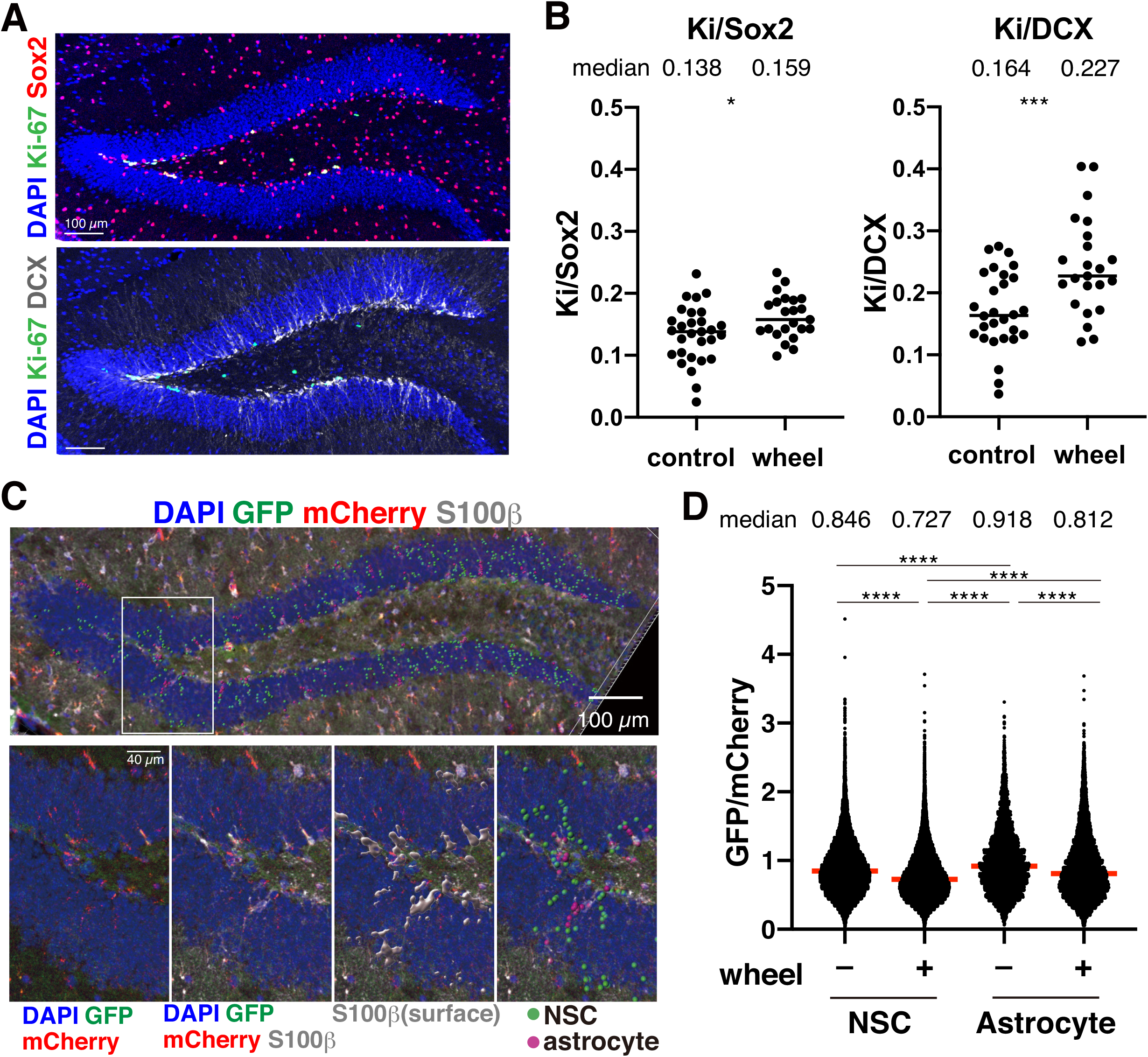
Lysosomal activity in NSCs of the DG of mice with or without running wheel. **A. Measurement of proliferating NSCs and newly-born neurons**. Representative photos for counting proliferating Ki-67+ (green)/Sox2+ (red) cells and Ki-67+ (green)/DCX+ (grey) cells in the DG. Ki-67+/Sox2+ cells are white in the upper panel. **B. Increased number of proliferating cells in the DG of mice with the wheel in cages**. The ratio of Ki-67+/Sox2+ to Sox2+ cells (left) and Ki-67+/DCX+ to DCX+ cells were calculated and plotted. Each data point represents results from one brain section. Bars represent median values. (\**P* < 0.05, \*\*\**P* < 0.001; Student’s t-test). Median values are above the chart. **C. Lysosprobe measurement in NSCs of the hippocampal DG**. Spots correspond mCherry (red)-positive lysosomes in the molecular layer of the DG (DAPI, blue). The grey surface is the S100β positive area. Magenta and green spots are mCherry of LyMo in S100β positive (astrocyte) and negative cells (NSC), respectively. **D. Activation of lysosomal protein degradation in the DG**. GFP and mCherry ratio was measured in spots (C) and adjusted with the median of the control sample (2M wild type in Fig. 4E). Red bars represent medians. (\*\*\*\**P* < 0.0001; one-way ANOVA with Tukey’s multiple comparison tests, n=6). Median values are above the chart.

### Lysosomal activity fluctuation depends on maturation, age, and disease *in vivo*

Previous reports have shown that lysosomes contribute to the differentiation, quiescence, and aging of NSCs (Kobayashi *et al*., 2019; Leeman *et al*., 2018; Yuizumi *et al*., 2021). In addition, it has recently been reported that lysosomal dysfunction with lowered v-ATPase activity in neurons precedes the onset of neurodegeneration associated with Alzheimer’s disease (Lee *et al*., 2022). Therefore, we next investigated changes in lysosomal activity in NSCs upon brain maturation, aging, and disease. First, we analyzed the activity state of NSCs in proliferation and neurogenesis in the brain of mice from juvenile (two weeks old) to old age (two years old). 34 % of Sox2-positive NSCs were double-positive for Ki-67 at two weeks of age but markedly decreased with age in the hippocampal DG; a similar decrease was observed in 5xFAD mice (Fig. 4A). The production of DCX-positive immature neurons was reduced in two-month-old 5xFAD mice compared to control healthy mice (9.2 % in controls and 7.3 % in 5xFAD mice; *p-value* < 0.0001), but was not significantly different at six months nor at one year (Fig. 4B). These results suggest that age-dependent reductions of both proliferation and neurogenesis have more dominantly appeared than disease-dependent ones in this time series. Neurogenesis was mainly affected by age in the adult brain in these transgenic mice.

**Figure 4.**
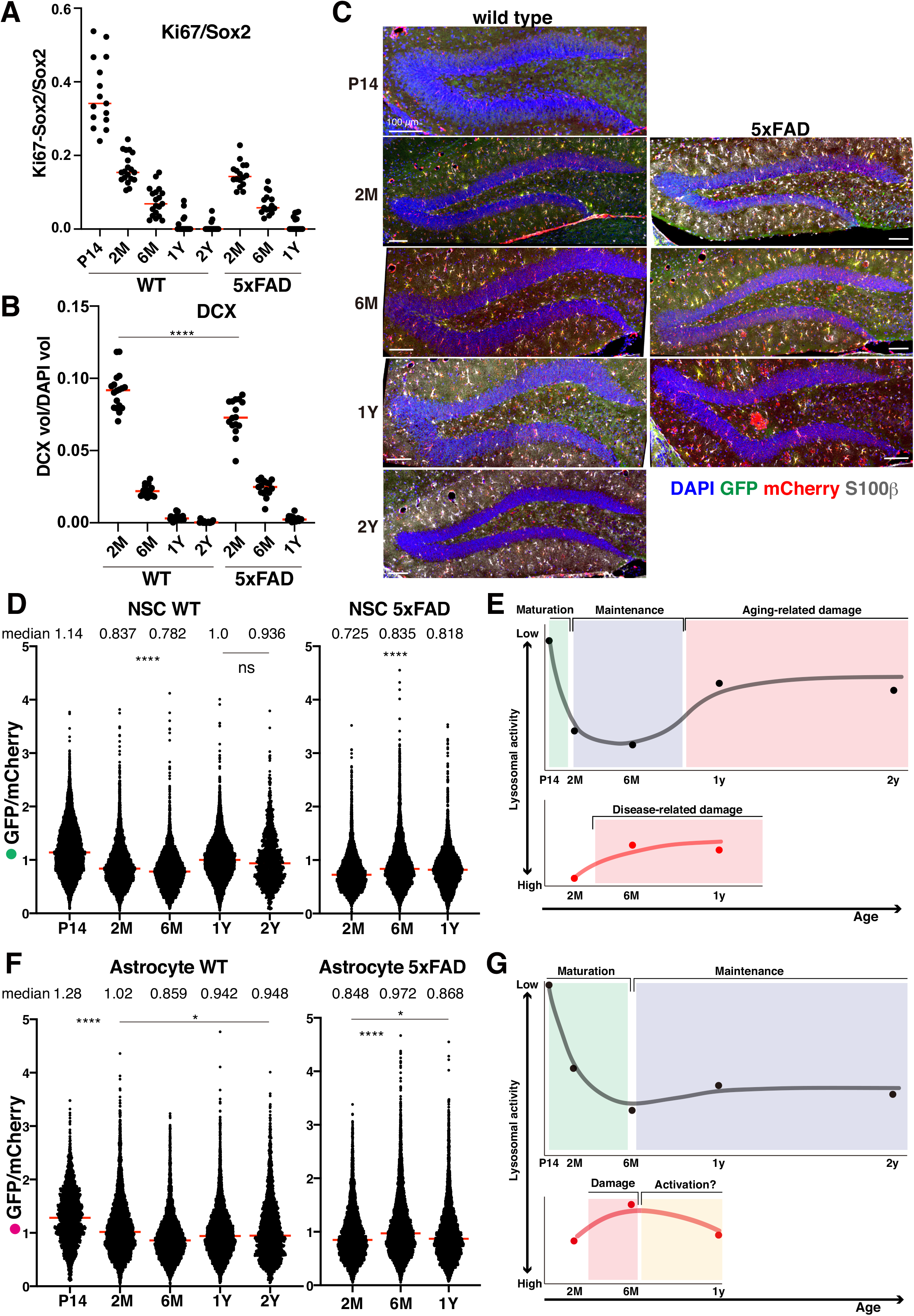
Lysosomal activity fluctuation in NSCs of the DG with age. **A, B. Measurement of proliferating NSCs and early-born neurons in the DG**. The ratio of Ki-67+/Sox2+ cells in all Sox2+ cells in the SGZ of the DG (A). DCX-positive volume was divided by the DAPI-positive volume (B). Each data point represents results from one brain section. Red bars represent median values. **C. LyMo wild-type and 5xFAD mice of different ages**. Representative images for measuring the lysosomal activity of different age mice and Alzheimer’s disease model mice. Scale bars, 100 μm. **D-G. Lysosomal protein degradation activities in NSCs (D) and astrocytes (F) of wild-type and 5xFAD mice and the hypothesized models of different lysosomal activity in NSCs (E) and astrocytes (G)**. GFP/mCherry ratio was measured and normalized with the median of the control NSCs (wild-type 1Y). Red bars represent medians (D, F). All compared pairs showed \*\*\*\**P < 0*.*0001* except some pairs described ns (not significant) or \**P<0*.*05*, and median values are above the chart (D, F). Dots in the charts show median values in the wild-type (black lines) and 5xFAD mice (red lines), respectively (E, G).

Next, we quantified lysosomal activity using LyMo in hippocampal NSCs in these mice (Fig. 4C). Lysosomal activity increased from age two weeks (P14) to two months (2M), increased slightly from age two months to six months, and decreased from age one year to two years (Fig. 4D). In 5xFAD mice, a decrease in lysosomal activity was observed between two and six months of age (Fig. 4D). These results demonstrated that lysosomal activity increased with brain maturation and was inversely correlated with the proliferation and differentiation potentials of NSCs from juvenile (two-weeks old) to adult periods (2 and 6 months old). However, this relationship was disrupted by further aging or disease: lysosomal activity suddenly reduced in NSCs at 1 and 2 years in control wild-type mice and reduced earlier, between 2 and 6 months old, in 5xFAD mice than in the control mice (Fig. 4E). On the other hand, in astrocytes of the hippocampal DG in control non-disease mice, the lysosomal activity increased from two weeks to six months of age, similar to NSCs, and then decreased after six months to slightly less in astrocyte than that in NSCs (Fig. 4F, G). This result suggests that the lysosomal activity was still somewhat maintained in astrocytes in aged mouse brains. However, in 5xFAD mice, lysosomal activity transiently reduced at six months and recovered at one year of age. This change was puzzling as higher lysosomal activity at an older age (2-year-old), but the findings may suggest that disease-dependent damage, for instance, the accumulation of amyloid-β plaques, reduces lysosomal activity at first and then later, lysosomal activity increases in response to this damage via astrocyte activation.

Our approach to measuring *in vivo* lysosomal activity using LyMo mice revealed that the fluctuation of lysosomal activity in NSCs depends on brain maturation, age, disease stage, and physical exercise. Lysosomal activity has been detected by quenching of GFP fluorescence due to lower pH in lysosomes; however, our new probe, LyMo, enabled us to monitor the “protein degradation” of destabilized GFP in lysosomes. Moreover, our method enables the measurement of lysosomal activity in NSCs of the hippocampal DG with spatial information and characterization by IHC. This is the first report to monitor the fluctuation of lysosomal protein degradation activity in NSCs *in vivo*. Still, it was technically challenging to compare lysosomal activity in individual cells because lysosomes are located throughout NSCs – in the soma, axon, and dendrites. Our measurements suggest that lysosomal activity might be helpful as an early and sensitive indicator of cellular condition in the brain along life stages. This method would likely apply to other tissue to measure lysosomal activity and cellular state *in vivo*.

## Methods

### Cell culture

NSCs were grown in DMEM/F-12 (Gibco) supplemented with 20 ng/ml EGF (Wako), 20 ng/ml bFGF (Wako), P/S (Nacalai Tesque), and N-2 max media supplement (R&D) (Kobayashi *et al*., 2019). NSCs expressing LyMo were generated by lentiviral transduction coding a Tet-On inducible cassette (Ishii *et al*., 2019; Kobayashi *et al*., 2019) and incubated for one day in the presence of doxycycline. Inhibitors were incubated at the following concentration: 20 nM bafilomycin A1 (Sigma), 100 nM Trin 1 (Cayman Chemical), and 10 μg/ml cycloheximide (Sigma). For immunocytochemistry, cells were fixed in 4% PFA/PBS on ice, permeabilized with 0.1% Triton X-100 in PBS (PBST), blocked in 5% normal goat serum/PBST and stained with antibodies in 1% normal goat serum/PBST. For western blotting, cells were lysed in lysis buffer (50 mM Tris-HCl (pH 8.0), 100 mM NaCl, 5 mM MgCl2, 0.5% Nonidet P-40, protease inhibitor cocktail (Roche), 1mM phenylmethylsulfonyl fluoride, 250 U/ml Benzonase (Sigma), and phosphatase inhibitors on ice after cold PBS wash, and subjected to SDS-PAGE.

### Mice

Mice were maintained in our animal facility and housed in a 12:12 hour light–dark cycle. Animal care and experiments were conducted per the guidelines of the Kyoto University animal experiment committee. In addition, we have complied with all relevant ethical animal testing and research regulations. LyMo Tg mice were created by injecting fertilized eggs of the C57BL/6 mice with DNA fragments containing the mouse GFAP promoter, lysosomal probe-coding sequences, and rabbit β-globin polyadenylation signal. For voluntary running experiments, the Tg mice were individually housed in cages with or without a running wheel at six weeks old and sacrificed after six weeks. In addition, 5XFAD Tg mice were obtained from Jackson Lab (#034840-JAX B6SJL-Tg (APPSwFlLon, PSEN1*M146L*L286V) 6799Vas/Mmjax) and crossed with LyMo mice to make double Tg mice.

### Brain sample preparation

For immunohistochemistry of the brain’s dentate gyrus (DG), mice were transcardially perfused with 4% PFA/PBS. Brains were postfixed with 4% PFA/PBS, cryoprotected with sucrose/PBS, embedded and frozen in OCT compound (Tissue TEK), and cryosectioned with 20 μm thickness. Every twelfth slice of 20 μm cryosections taken along the caudal-rostral axis throughout the entire DG was immunostained for quantification. For slice culture, brains were dropped into cutting solution (280 mM sucrose, 2 mM KCl, 10 mM HEPES-NaOH [pH 7.4], 0.5 mM CaCl2, 10 mM MgCl2, and 10 mM glucose) and sliced into 200 μm-thick slices on a vibratome (Leica, VT1200S). Brain slices were incubated in bath solution (135 mM NaCl, 5 mM KCl, 1 mM CaCl2, 1 mM MgCl2, 10 mM HEPES-NaOH [pH 7.4] and 10 mM glucose) for 30 min with bubbling oxygen and then transferred on Millicell Cell Culture Inserts (0.4-μm pore size, EMD Millipore) covered with Cellmatrix Type I-A collagen gel (Nitta Gelatin) and incubated overnight in the culture medium (10% FBS/ bath solution) in absence or presence of bafilomycin A1. Slices were fixed with 4% PFA/PBS for one hour and immunostained after removal from the collagen gel. Six slices per condition were measured.

### Antibodies

Chicken anti-GFP (Abcam, ab13970), goat anti-mCherry (SICGEN, AB0040), rat anti-Lamp1 (Developmental Studies Hybridoma Bank, 1D4B), mouse anti-GFAP (sigma, G3893), goat anti-Sox2 (R&D Systems, AF2018), chicken anti-Nestin (aves, NES), rabbit anti-S100β (Abcam, ab52642), rat anti-Ki-67(eBioscience, SolA15), and rabbit anti-DCX (CST, 4604) antibodies were used for immunohisto- and immunocytochemistry. Rabbit anti-GFP (Thermo, A11122), rabbit anti-mCherry (Abcam, ab167453), and rabbit anti-actin (Sigma, A2066) antibodies were used for western blotting. In addition, we used donkey or goat antibodies conjugated with Alexa 488, 568, or 647 (Thermo, Abcam) and goat antibodies conjugated with HRP (GE) as secondary antibodies.

### IHC

Brain cryosections were incubated in Histo-VT one (Nakalai) at 70 °C for 20 min, permeabilized and blocked with 5% normal donkey serum (NDS)/0.1% Triton X-100/PBS (PBST), and stained with 1^st^ and 2^nd^ antibodies with DAPI (Sigma) in PBST. Aged brain samples (2Y) were treated with an autofluorescence eliminator reagent (Millipore) to reduce lipofuscin-related background. Brain slices after slice culture were permeabilized with 0.3% Triton X-100/PBS before blocking with 5% normal donkey serum (NDS)/0.3% Triton X-100/PBS and stained with 1^st^ and 2^nd^ antibodies in 1% NDS/0.3% Triton X-100/PBS. Confocal z-stack images were obtained on Stellaris 5 (Leica) and LSM880 Airyscan (Zeiss).

### Counting proliferating NSCs and newly born neurons

Imaris software was used for all counting using 3D confocal images. Sox2, DCX, and Ki-67 were masked by the DAPI surface (5 μm surface detail) to extract cells in the granular cell layer of the DG. Spots were put on Sox2-positive (XY diameter 5μm, signals with both max intensity above 2500 and mean intensity above 700) and DCX-positive (XY diameter 10 μm, signals with quality above 1000) areas for counting. Ki-67 intensity (intensity standard deviation above 2700) was used to filter the spots for counting Ki-67-Sox2 and Ki-67-DCX double-positive cells.

### Quantification of lysosomal activity by LyMo

In cells after immunocytochemistry, spots for measurement were picked up as mCherry-positive dots of lysosomes with 0.8 μm diameter and 6000 quality value in Imaris software. mCherry and GFP intensity on these spots were measured, and GFP intensity was divided by mCherry intensity and normalized with the median value of the control sample. In the western blotting of NSCs, both LyMo full-length and GFP-mCherry fusion was detected by both GFP and mCherry antibodies at 100 kDa and 50 kDa. The bands of GFP alone, mCherry alone, and actin were quantified on a LAS3000 image analyzer (Fujifilm) and normalized by the corresponding intensity of β-actin. In brain tissues after immunohistochemistry, the DAPI surface covering the granular zone masked mCherry and S100β signals and spots were placed on mCherry-positive lysosomal signals with 2 μm diameter. mCherry spots were filtered by “quality” and the top 2 % of spots were analyzed. To isolate S100β-positive astrocytes, an additional S100β surface was made within the DAPI-mask using S100β signals (1.14 μm surface detail), distinguishing mCherry spots inside and outside the distance value from S100β surface. GFP intensity / mCherry intensity ratio was calculated with Python.

### Statistical analysis

Statistical analyses were performed using GraphPad Prism8 software. Statistical differences were examined using Student’s t-test, one-way ANOVA, and two-way ANOVA with Tukey’s multiple comparisons tests. P-values < 0.05 were considered significant.

## Acknowledgments

We thank F. Ishidate, Y. Kosodo, M. Matsuda, and members of the Kageyama laboratory and the Matsuda laboratory for technical help and discussion; iCems analysis center, innovative support alliance for life science (iSal), and research center for dynamic living systems in Kyoto University, for technical help. This work was supported by a Grant-in-Aid for Scientific Research (B) (JSPS 20H03260) (TK), Grant-in-Aid for Transformative Research Areas (JSPS 23H04919) (TK), AMED under Grant Number JP20gm6410006 (TK), and SPIRITS of Kyoto University (TK).

## Author contributions

HZ, KI, and TK conducted all of the experiments. TS performed data processing of results from LyMo mice. SI and EI established the construct of the LyMo probe. MH and ZL helped ZH with mouse experiments. RT established the NSC line expressing LyMo. SK and HM helped to obtain transgenic mice by *in vitro* fertilization. RK provided 5xFAD mice. TK designed the experiments and wrote the manuscript; KI edited the English.

## References

Ballabio A, Bonifacino JS (2020) Lysosomes as dynamic regulators of cell and organismal homeostasis. Nat Rev Mol Cell Biol 21: 101–118

Cole JD, Sarabia Del Castillo J, Gut G, Gonzalez-Bohorquez D, Pelkmans L, Jessberger S (2022) Characterization of the neurogenic niche in the aging dentate gyrus using iterative immunofluorescence imaging. Elife 11: e68000

Garrett L, Lie DC, Hrabe de Angelis M, Wurst W, Holter SM (2012) Voluntary wheel running in mice increases the rate of neurogenesis without affecting anxiety-related behaviour in single tests. BMC Neurosci 13: 61

Huang J, Wang X, Zhu Y, Li Z, Zhu YT, Wu JC, Qin ZH, Xiang M, Lin F (2019) Exercise activates lysosomal function in the brain through AMPK-SIRT1-TFEB pathway. CNS Neurosci Ther 25: 796–807

Ishii S, Matsuura A, Itakura E (2019) Identification of a factor controlling lysosomal homeostasis using a novel lysosomal trafficking probe. Sci Rep 9: 11635

Karpf J, Unichenko P, Chalmers N, Beyer F, Wittmann MT, Schneider J, Fidan E, Reis A, Beckervordersandforth J, Brandner S et al (2022) Dentate gyrus astrocytes exhibit layer-specific molecular, morphological and physiological features. Nat Neurosci 25: 1626–1638

Katayama H, Yamamoto A, Mizushima N, Yoshimori T, Miyawaki A (2008) GFP-like proteins stably accumulate in lysosomes. Cell Struct Funct 33: 1–12

Khmelinskii A, Keller PJ, Bartosik A, Meurer M, Barry JD, Mardin BR, Kaufmann A, Trautmann S, Wachsmuth M, Pereira G et al (2012) Tandem fluorescent protein timers for in vivo analysis of protein dynamics. Nat Biotechnol 30: 708–714

Kobayashi T, Piao W, Takamura T, Kori H, Miyachi H, Kitano S, Iwamoto Y, Yamada M, Imayoshi I, Shioda S et al (2019) Enhanced lysosomal degradation maintains the quiescent state of neural stem cells. Nat Commun 10: 5446

Kriegstein A, Alvarez-Buylla A (2009) The glial nature of embryonic and adult neural stem cells. Annu Rev Neurosci 32: 149–184

Lee JH, Yang DS, Goulbourne CN, Im E, Stavrides P, Pensalfini A, Chan H, Bouchet-Marquis C, Bleiwas C, Berg MJ et al (2022) Faulty autolysosome acidification in Alzheimer’s disease mouse models induces autophagic build-up of Abeta in neurons, yielding senile plaques. Nat Neurosci 25: 688–701

Leeman DS, Hebestreit K, Ruetz T, Webb AE, McKay A, Pollina EA, Dulken BW, Zhao X, Yeo RW, Ho TT et al (2018) Lysosome activation clears aggregates and enhances quiescent neural stem cell activation during aging. Science 359: 1277–1283

Leiter O, Seidemann S, Overall RW, Ramasz B, Rund N, Schallenberg S, Grinenko T, Wielockx B, Kempermann G, Walker TL (2019) Exercise-Induced Activated Platelets Increase Adult Hippocampal Precursor Proliferation and Promote Neuronal Differentiation. Stem Cell Reports 12: 667–679

Mizushima N, Murphy LO (2020) Autophagy Assays for Biological Discovery and Therapeutic Development. Trends Biochem Sci 45: 1080–1093

Morrow CS, Porter TJ, Xu N, Arndt ZP, Ako-Asare K, Heo HJ, Thompson EAN, Moore DL (2020) Vimentin Coordinates Protein Turnover at the Aggresome during Neural Stem Cell Quiescence Exit. Cell Stem Cell 26: 558–568 e559

Neefjes J, Dantuma NP (2004) Fluorescent probes for proteolysis: tools for drug discovery. Nat Rev Drug Discov 3: 58–69

Oakley H, Cole SL, Logan S, Maus E, Shao P, Craft J, Guillozet-Bongaarts A, Ohno M, Disterhoft J, Van Eldik L et al (2006) Intraneuronal beta-amyloid aggregates, neurodegeneration, and neuron loss in transgenic mice with five familial Alzheimer’s disease mutations: potential factors in amyloid plaque formation. J Neurosci 26: 10129–10140

Ohkouchi S, Shibata M, Sasaki M, Koike M, Safig P, Peters C, Nagata S, Uchiyama Y (2013) Biogenesis and proteolytic processing of lysosomal DNase II. PLoS One 8: e59148

Pedelacq JD, Cabantous S, Tran T, Terwilliger TC, Waldo GS (2006) Engineering and characterization of a superfolder green fluorescent protein. Nat Biotechnol 24: 79–88

Seki T, Sato T, Toda K, Osumi N, Imura T, Shioda S (2014) Distinctive population of Gfap-expressing neural progenitors arising around the dentate notch migrate and form the granule cell layer in the developing hippocampus. J Comp Neurol 522: 261–283

Settembre C, Zoncu R, Medina DL, Vetrini F, Erdin S, Erdin S, Huynh T, Ferron M, Karsenty G, Vellard MC et al (2012) A lysosome-to-nucleus signalling mechanism senses and regulates the lysosome via mTOR and TFEB. EMBO J 31: 1095–1108

van Praag H, Kempermann G, Gage FH (1999) Running increases cell proliferation and neurogenesis in the adult mouse dentate gyrus. Nat Neurosci 2: 266–270

Yanai S, Endo S (2021) Functional Aging in Male C57BL/6J Mice Across the Life-Span: A Systematic Behavioral Analysis of Motor, Emotional, and Memory Function to Define an Aging Phenotype. Front Aging Neurosci 13: 697621

Yuizumi N, Harada Y, Kuniya T, Sunabori T, Koike M, Wakabayashi M, Ishihama Y, Suzuki Y, Kawaguchi D, Gotoh Y (2021) Maintenance of neural stem-progenitor cells by the lysosomal biosynthesis regulators TFEB and TFE3 in the embryonic mouse telencephalon. Stem Cells 39: 929–944

